# The Diverse Phenotypic and Mutational Landscape Induced by Fluoroquinolone Treatment

**DOI:** 10.1101/2024.12.20.629600

**Authors:** Sayed Golam Mohiuddin, Pouria Kavousi, Diego Figueroa, Sreyashi Ghosh, Mehmet A. Orman

## Abstract

Despite extensive research on antibiotic resistance, the potential effects of antibiotic treatments on bacterial tolerance and resistance remain a significant concern. Although bacterial cells adopt a variety of mutational strategies to resist unfavorable circumstances, it is still unclear how antibiotic tolerance and resistance mechanisms affect bacterial fitness characteristics and whether evolved mutants exhibit similar properties across different cell populations subjected to the same conditions. Here, we used *Escherichia coli*, a fluoroquinolone antibiotic (ofloxacin), and adaptive laboratory evolutionary experiments to demonstrate that ofloxacin tolerance and resistance can evolve independently across different cell populations exposed to identical conditions. Fitness attributes, such as lag score, doubling time, competition score, and other metabolic features, were variably affected by antibiotic tolerance and resistance mechanisms. However, we did not observe strong and apparent correlations between fitness trade-offs and antibiotic tolerance and resistance. While our whole-genome sequencing identified some shared mutations, such as single nucleotide polymorphisms in the *icd* gene (a crucial citric acid cycle gene), evolved cell populations exhibited diverse genetic mutations without a clear pattern of a conserved evolutionary pathway. Our study also identifies unique phenotypes, such as those displaying significantly lower minimum inhibitory concentration levels compared to the parental strain yet showing remarkably high tolerance to the same antibiotic. Altogether, our study, examining the phenotypic and mutational landscapes of fluoroquinolone-induced strains, contributes to our understanding of complex bacterial adaptation mechanisms.

## Introduction

Referred to as the hidden pandemic [1], the evolution of resistant mutations that render antibiotic therapy ineffective has been identified as a serious global public health concern [2,3]. According to a “United Nations” report, antibiotic resistance will kill about 10 million people by 2050, which will potentially match the number of fatalities from cancer [4,5]. A reduction in treatment options could push public health into a post-antibiotic era that can significantly increase the number of infectious disease cases [6]. Numerous mechanisms, including reversible and irreversible processes that bacteria can use to survive antibiotics, have been linked to the failure of antibiotic therapies. Bacterial persistence is a reversible survival mechanism that allows a small fraction of a genetically identical cell population to tolerate high doses of antibiotics transiently [7–11]. This differs from antibiotic resistance, which involves an increase in the minimum inhibitory concentration (MIC) of antibiotics in resistant cells, typically acquired through genetic mutations [12–15]. While resistance and persistence are two different phenomena, they might be related, given that persister cells can act as a reservoir for the formation of resistant mutations [16–22]. Under certain conditions, cells may acquire mutations that enhance survival at the population level without altering MIC levels; we will refer to this phenomenon as "tolerance" in this study to distinguish it from resistance.

Bacteria can acquire resistance through various means, including plasmid-mediated conjugation that facilitates the lateral transfer of resistance genes between bacteria and other microorganisms [23–26]. Furthermore, a variety of environmental factors, such as nutrient depletion, UV radiation, antibiotics, and toxic chemicals, can alter the genetic makeup of bacterial chromosomes and may lead to structural anomalies, deletions, insertions, and single nucleotide polymorphisms that can make the cells tolerant and/or resistant [16,17,27–30]. In particular, the extensive use of antibiotics that damage DNA, like fluoroquinolones, can significantly increase bacterial mutagenesis by activating error-prone DNA repair systems [17,31–37]. Mutations that result in antibiotic resistance may modify drug target sites, efflux pumps, membrane structure, metabolic pathways that may enhance cell dormancy, and enzymes that deactivate antibiotics [38–41]. Although antibiotic resistance has been extensively studied [42–45], the potential impact of antibiotic treatment on bacterial resistance and persistence is a matter of great concern, underscoring the need for further research in this area.

Adaptive laboratory evolutionary (ALE) experiments provide critical insights into the mechanisms by which bacteria adapt and develop resistance in response to antibiotic exposure [16,17,19,21,46]. For instance, *in vitro* controlled experimental evolution during ampicillin treatment demonstrated that tolerance paves the way for the emergence of antibiotic-resistant mutants, and a mathematical model describing population genetics showed how tolerance accelerates the rate of resistant mutations in a subpopulation [16]. Experimental data in combination with the hypothetical model demonstrated that persistence is pleiotropically linked to increased mutation rates and consequently acts as the source of antibiotic-resistant mutants [16]. Using classical and metabolism-dependent antibiotic-mediated evolutionary experiments demonstrated the presence of mutations in the metabolic genes that were shown to be clinically relevant [21].

ALE experiments, persistence, and mutagenesis are intricately linked to bacterial adaptation and antibiotic resistance. During ALE experiments, persister cells—variants that survive initial treatments—play a crucial role as a reservoir for survival [16,17,19,21]. Although not inherently resistant, these cells can undergo mutagenesis, especially when exposed to stressors like antibiotics that induce DNA damage and error-prone repair mechanisms [17,19,21,47]. The cyclic nature of ALE, involving alternating phases of antibiotic exposure and recovery, further amplifies this process by allowing surviving cells to proliferate and compete, potentially acquiring and fixing beneficial mutations that confer resistance. Unlike traditional models, which suggest that resistance mutations typically come with fitness costs such as slower growth rates [16,48–51], the dynamic environment of cyclic ALE can favor the selection of mutations that confer resistance while maintaining overall fitness. These adaptive responses to fluctuating environments, commonly observed in real-life clinical and environmental settings [52,53], underscore the need for further studies to clarify how these processes impact bacterial fitness.

Although the convergence of evolutionary trajectories towards a common pathway or mechanism is highly sought after by the scientific community [16,17,19,21,54] —a crucial aspect for identifying therapeutic targets—, living organisms have evolved over billions of years, resulting in complex genetic landscapes. This diversity has given them a multitude of mechanisms to survive adverse conditions. In this study, we aimed to address several scientific questions in the field using ALE experiments. These questions included understanding the survival and resistance traits of evolved bacteria under antibiotic pressure, assessing whether evolved mutants exhibit consistent phenotypic traits within and across different samples, exploring how antibiotic tolerance and resistance affect other fitness attributes of bacteria, investigating the genetic mutations contributing to observed tolerance and resistance, and examining whether there is a conserved mechanism that could be explored for therapeutic purposes. By tackling each of the aforementioned issues, our research advances our knowledge of the mechanisms underlying bacterial adaptability and its heterogeneous nature.

## Results

### Adaptive laboratory evolutionary experiments generated diverse tolerant and resistant strains

We used an *in vitro* ALE methodology to investigate whether sporadic antibiotic exposure could yield mutants displaying increased tolerance or resistance. To carry out this experiment, we utilized an *Escherichia coli* MG1655 strain (MO-cured [55]) harboring both a chromosomally integrated inducible mCherry expression system and a pUA66-empty vector (EV) with kanamycin resistance gene (Supplementary **Table S1**). Kanamycin (50 µg/mL) was used to maintain the plasmid and prevent contamination, given the prolonged duration of the experiment. We used mCherry-expressing cells, which proved highly useful in analyzing the fitness characteristics of the generated mutant strains. We initiated the ALE experiment by preparing eight separate 2 mL-overnight cultures from the same cell stock, incubating them under identical conditions in a shaker at 37°C and 250 rpm for 16 hours. These cultures were then diluted in 2 mL of fresh Lysogeny Broth (LB) medium and promptly exposed to a high concentration of ofloxacin (5 µg/mL) for a duration of 7 hours. Subsequently, the treated cultures were collected and washed with phosphate-buffered solution (PBS) to remove antibiotics, and half of the washed cells were plated to quantify the surviving cells. The remaining cells were inoculated in 2 mL of fresh LB medium and allowed to grow overnight (16 h) with shaking at 37° C to facilitate the recovery of the surviving cells, and this cycle was repeated over a span of 22 days. To visualize the trajectory of survival, cells were plated on LB agar plates both before and after the ofloxacin treatment in each cycle of the ALE experiment (**Fig. 1a**). Throughout the course of the experiment, the trajectories of eight distinct samples exhibited varying forms of adaptation to the high concentration of ofloxacin. <u>S</u>ample <u>2</u> (S2) displayed no discernible increased survival to ofloxacin even after intermittent exposure for 22 days (**Fig. 1a**), akin to the parental strain (without 22 days of ALE). Conversely, seven samples (S1, S3, S4, S5, S6, S7, and S8) exhibited an increased survival to ofloxacin (**Fig. 1a**) compared to the control (parental strain).

**Fig. 1.**
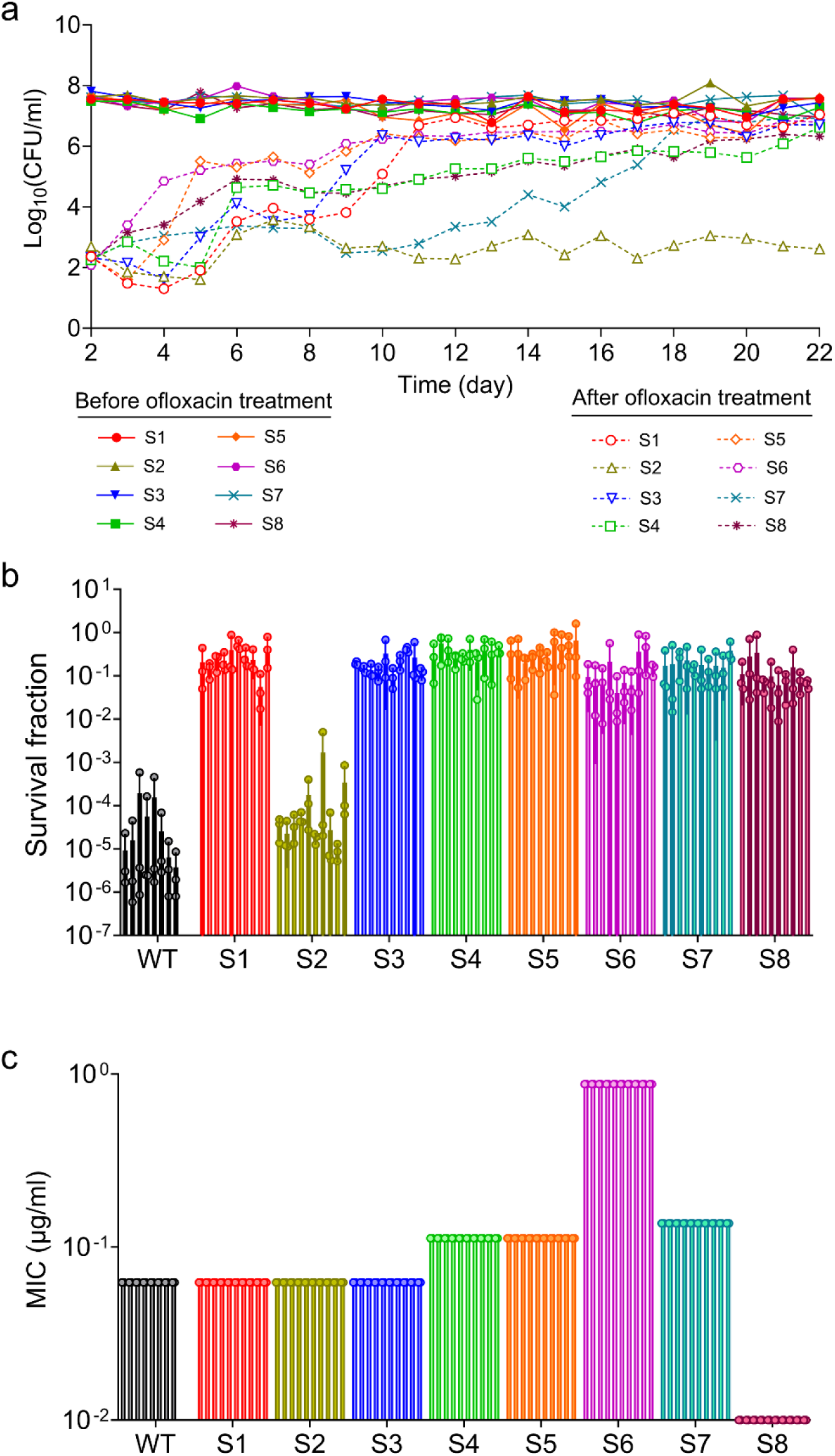
Generation of ofloxacin mutant strains using adaptive laboratory evolutionary experiment. **(a)** Stationary phase cells (16 h overnight culture) of *E. coli* MG1655 MO strain harboring pUA66-empty vector plasmid, were exposed to ofloxacin (5 µg/ml) after diluting 100- fold in LB media for 7 h. Treated cells were collected and washed to remove the antibiotics. Washed cells (1 ml) were then transferred to fresh LB media and grew overnight. The remaining 1 ml washed cells were plated on LB agar plate to enumerate the colony formation unit (CFU). These steps were repeated for 22 days. CFU levels before and after the ofloxacin treatment have been reported in the plot. Number of independent biological replicates, n = 1. **(b)** Stationary phase cells of ten randomly picked colonies of the indicated strains were diluted 100-fold in LB and treated with ofloxacin (5 µg/ml) for 7 h. After the treatment, cells were washed and plated on LB agar to determine the survival fractions. n=3. **(c)** Ten colonies of the indicated strains were diluted to have ∼10^8^-10^9^ cells/ml and spread on LB agar plates. Ofloxacin ETEST strips were placed on the plates, which were then incubated at 37°C for 16 hours. The parabola formed after incubation was used to determine the minimum inhibitory concentrations. n=3. Data corresponding to each time point represents mean value ± standard deviation.

Given the random nature of mutations, each of the 8 samples may harbor distinct and numerous ofloxacin mutants. To determine whether these mutants display consistent phenotypic traits within each sample, we streaked cells from each sample onto LB agar plates. We then randomly selected at least ten individual colonies from each sample to assess their tolerance and/or resistance properties by measuring ofloxacin survival fractions and minimum inhibitory concentrations (MIC). Although there is noteworthy variation in survival fractions among the 8 samples, as depicted in **Fig. 1a**, the survival fractions among the ten colonies within each sample were found to be similar in **Fig. 1b**. Our findings also revealed consistent MIC levels among the ten selected colonies within each sample, while there were notable variations in MIC levels across the 8 samples, as demonstrated in **Fig. 1c**. Surprisingly, S1, S2, and S3 displayed MIC levels identical to those of the parental strains (**Fig. 1c**). On the other hand, the ofloxacin MIC level of sample S8 was significantly lower than that of the parental strain despite its markedly higher survival fractions (**Fig. 1b, c**). This trend differs in sample S7, as we observed both higher MIC levels and increased survival to ofloxacin compared to the parental strain. (**Fig. 1b, c**). However, a correlation analysis between all eight samples’ survival fractions and MIC levels (Supplementary **Fig. S1**) demonstrated that these two phenomena are not always related (R^2^ = 0.015, P = 0.48, F-statistics).

Altogether, these experiments yielded two key findings:

i. In each sample, all ten random colonies selected have the same MIC and survival fractions, as depicted in **Fig. 1b** and **1c**, highlighting the enrichment of mutant strains with consistent phenotypic characteristics within each sample. Surprisingly, these traits were not conserved across the eight different samples, even though they were generated under identical conditions.
ii. It was evident that survival fractions and MIC levels did not always correlate. We have identified strains exhibiting significantly lower MIC levels than the parental strain while simultaneously displaying remarkably high tolerance to the same antibiotic (**Fig. 1b** and **1c**). To the best of our knowledge, these strains have not been comprehensively studied or documented previously.

### Antibiotic-tolerant cells exhibit diverse fitness attributes

A trade-off can exist between antibiotic tolerance/resistance and other traits. For example, mutations that confer antibiotic resistance may come with associated costs, potentially reducing the bacterium’s overall fitness. To explore the connections between the tolerance/resistance of our samples and their fitness, we evaluated various fitness indicators, such as the lag score, doubling time, competition score, non-growing score, cellular redox activities, and ATP levels.

The lag score, representing the duration that bacteria take to initiate growth in a specific medium, was determined using the methodology outlined in a previous study [56,57]. It was evident that samples S4 and S8 exhibited a notable increase in lag scores compared to the control (**Fig. 2a**, Supplementary **Fig. S2a**). Conversely, samples S2, S3, S5, and S6 displayed lag scores like the control (**Fig. 2a**, Supplementary **Fig. S2a**). Our attempt to correlate survival fraction data with the lag score of each sample was motivated by existing literature, which suggests that cell tolerance to antibiotics is influenced by the length of the lag phase—where a longer lag phase corresponds to higher antibiotic tolerance [16,48]. Our findings revealed no clear correlation between the lag scores and survival fractions of all eight samples (R^2^ = 0.16, P = 0.016, F-statistics, **Fig. 3a**). Another parameter potentially contributing to the higher survival fraction of the ALE-evolved strains is doubling time. Apart from sample S2, all other evolved samples exhibited significantly higher doubling times compared to the control strain (**Fig. 2a**, Supplementary **Fig. S2b**). Unlike lag scores, when comparing survival fractions and doubling times for all samples, we found a moderate correlation between these two parameters (R^2^ = 0.71, P < 0.0001, F-statistics, **Fig. 3b**), indicating that doubling time may be a potential indicator of enhanced tolerance or resistance.

**Fig. 2.**
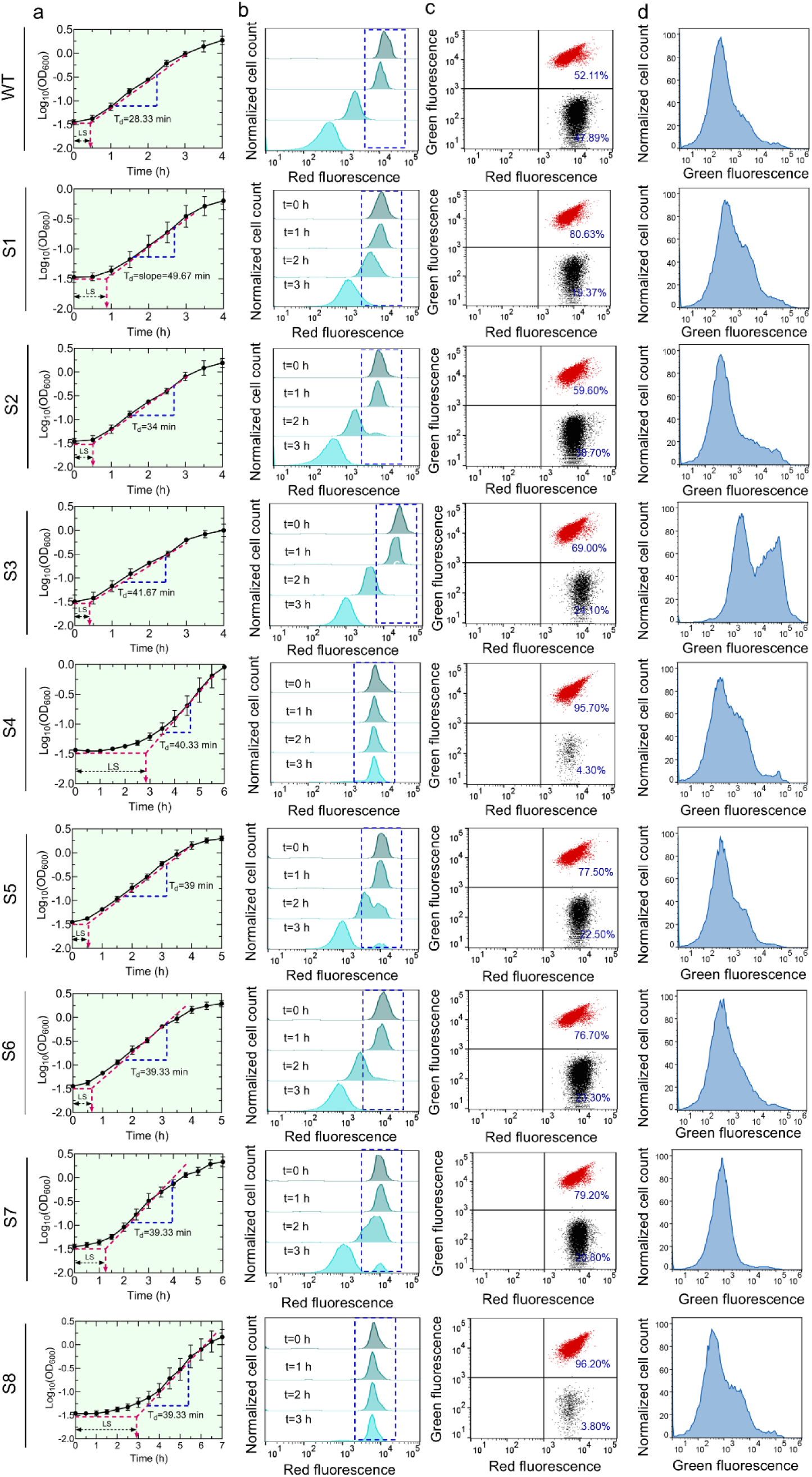
Determination of fitness factors and metabolic activities of the evolved strains. **(a)** Growth curves of the mutants and WT strain (control). Stationary phase cells were diluted 100-fold in LB media and cultured for 24 h. At designated time points, cells were collected to measure OD_600_ using a plate reader. These growth curves were utilized to determine the lag scores and doubling times (for details see Materials and Methods). n=4. **(b)** Flow-cytometry-based approach for the quantification of non-growing cell levels. The mutants and WT cells (harboring an IPTG inducible mCherry expression cassette in their genome) were cultured overnight in the presence of 1 mM IPTG to express the mCherry protein. Stationary phase cells were collected and washed to remove the IPTG and diluted 100-fold in LB media and grown in the shaker in the absence of IPTG. Cell division along with protein dilution were monitored using a flow-cytometer at single-cell levels at indicated time points. Non-growing cells retained their mCherry levels. A representative biological replicate is shown, with all three biological replicates consistently yielding similar trends. n=3. **(c)** Competition assays for mutants and WT strains in cocultures. Mutants (harboring pUA66-EV) and WT cells (harboring pUA66-*gfp*) were cultured individually overnight (16 h) in the presence of 1 mM IPTG in LB media. Stationary phase cells of the mutants and WT cells were diluted 100-fold in LB and cocultured in the presence of 1 mM IPTG for 24 h. At t=24, cells were collected, diluted in PBS and analyzed with a flow cytometer at the single-cell level, showing two distinct cell populations. For WT: red dots represent WT cells carrying both mCherry and GFP expression systems, while black dots represent WT cells carrying only the mCherry expression system. For samples S1-S8: red dots represent WT cells carrying both mCherry and GFP expression systems, while black dots represent the mutant cells carrying only the mCherry expression system. A representative biological replicate is shown, with all three biological replicates consistently yielding similar trends. n=3. **(d)** RSG staining of stationary phase cells. Stationary phase mutants and WT cells were stained with Redox Sensor Green (RSG) dye and analyzed with a flow cytometer to measure their metabolic activities. A representative flow cytometry diagram is shown. All independent biological replicates showed a similar trend. n = 3. Data corresponding to each time point represents mean value ± standard deviation.

**Fig. 3.**
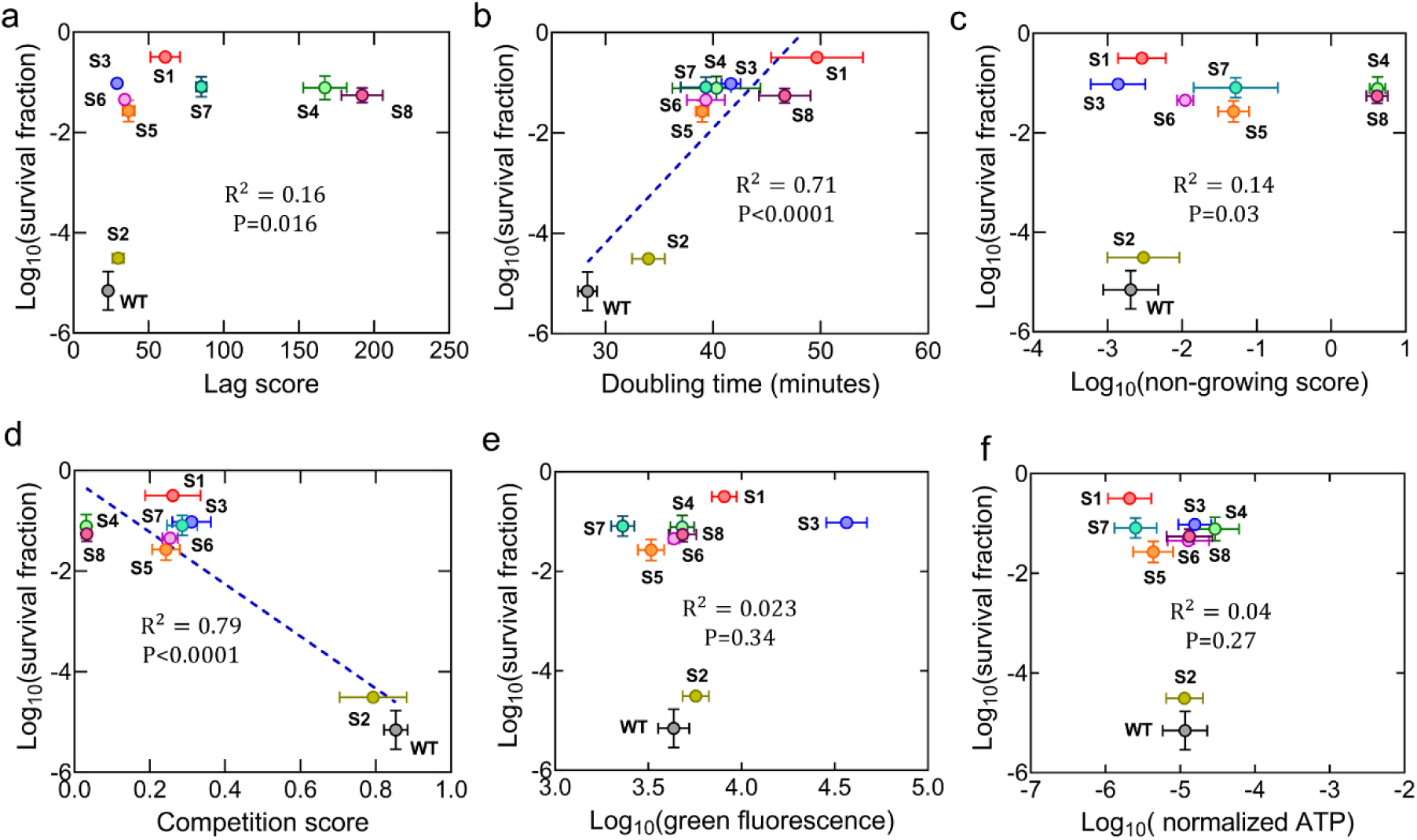
Correlation between survival fractions, fitness factors, and metabolic parameters of mutant strains. (a) Correlation between survival fractions and lag scores. A linear regression analysis was performed between survival fractions and lag scores of the mutant and WT strains (data from **Fig. 2a** and Supplementary **Fig. S2a**). (b) Correlation between survival fractions and doubling times. A linear regression analysis was performed between survival fractions and doubling times of the mutant and WT strains (data from **Fig. 2a** and Supplementary **Fig. S2b**). (c) Correlation between survival fractions and non-growing scores. A linear regression analysis was performed between survival fractions and non-growing scores of the mutant and WT strains (data from **Fig. 2b** and Supplementary **Fig. S2c**). (d) Correlation between survival fractions and competition scores. A linear regression analysis was performed between survival fractions and competition scores of the mutant and WT strains (data from **Fig. 2c** and Supplementary **Fig. S2d**). (e) Correlation between survival fractions and cellular redox state. A linear regression analysis was performed between survival fractions and metabolic states of the mutant and WT strains (data from Supplementary **Fig. S2e**). (f) Correlation between survival fractions and ATP levels. A linear regression analysis was performed between survival fractions and ATP levels of the mutant and WT strains (data from Supplementary **Fig. S2f**). Statistical significance was determined using F-statistics. Data corresponding to each time point represents mean value ± standard deviation.

The non-growing score, another significant parameter in defining fitness, was determined as the fraction of cells that did not undergo the cell division process at a specific time compared to the levels of actively growing cells. To determine the non-growing cell levels, we employed a fluorescent protein-dilution technique that we commonly utilized for monitoring cell growth [55,58,59]. In this process, we initially induced mCherry expression during the overnight growth of an *E. coli* strain containing a chromosomally integrated IPTG-inducible mCherry expression cassette [55,58]. The mCherry-positive cells from overnight cultures were then diluted in fresh medium and cultured without the inducer. Initially, all cells exhibited high levels of mCherry protein which gradually decreased with cellular growth due to protein dilution. However, a specific subpopulation still maintained elevated levels of red fluorescence owing to their lack of division. At t=3 h of growth, we assessed the levels of non-growing cells in each evolved sample and found that S4 and S8 exhibited significantly delayed growth resumption compared to the control strain (**Fig. 2b**, Supplementary **Fig. S2c**), which is expected as they had higher lag scores (**Fig. 2a**). In contrast, samples S1, S2, S3, S5, and S6 showed little or no non-growing cells, similar to the parental strain, while S7 had significantly more non-growing cells than the parental strain (**Fig. 2b**, Supplementary **Fig. S2c**). No definitive correlation emerged when we examined the correlation between non-growing scores and survival fractions (R^2^ = 0.14, P =0.03, F-statistics, **Fig. 3c**).

Competition scores represent the ratio of the mutant strain to the parental strain at a specific growth phase when they are cultured together. Our evolved strains contain an IPTG-inducible mCherry protein expression cassette in their genomic DNA and an empty plasmid (pUA66-EV). The parental strain, which also carries the mCherry expression system, contains pUA66-*gfp* plasmids encoding the green fluorescent protein (GFP). Cocultures were prepared by diluting and combining both the evolved and parental cells from separate overnight cultures into 2 mL fresh LB media in equal numbers (determined by a flow cytometer). Following 24 hours of culturing, the cocultures were analyzed using a flow cytometer to quantify mutant cells (mCherry only) and parental cells (mCherry+GFP), enabling the calculation of competition scores (**Fig. 2c**, see Supplementary **Fig. S3** for time-dependent flow cytometry data). We also utilized a parental strain with the empty plasmid as a control (black dots in the flow diagram for WT in **Fig. 2c**) to assess the impact of GFP overexpression on the competition score. While almost all mutant strains exhibited slower growth in co-cultures compared to the parental strain, S4 and S8 displayed significant growth inhibition, suggesting their inability to compete with parental cells (**Fig. 2c**, Supplementary **Fig. S2d**). Upon comparing survival fractions and competition scores for all samples, we found a moderate correlation between these two parameters (R^2^ = 0.79, P < 0.0001, F-statistics, **Fig. 3d**), indicating that the ability of a strain to survive and its competitive advantage in coculture settings may be interconnected. This is consistent with the correlation analysis of doubling time scores (**Fig. 3b**), as fast-growing strains are expected to compete better.

We evaluated the metabolic activities of mutant strains with Redox Sensor Green (RSG) dye and intracellular ATP measurements. RSG dye, capable of entering live bacteria, emits a green fluorescence signal when it undergoes reduction by bacterial reductase enzymes. RSG and ATP measurements were conducted immediately after diluting the overnight cultures, consistent with the ofloxacin treatment conditions. Interestingly, while the redox activities of most mutant strains were very similar to those of the parental cells, a significant increase in the redox activity of S3 was observed compared to that of the parental strain (**Fig. 2d**, Supplementary **Fig. S2e**). On the other hand, the S4 strain exhibited slightly higher ATP levels while S1, S5, and S7 demonstrated considerably lower ATP levels compared to the parental strain, though these differences were not statistically significant (Supplementary **Fig. S2f**). We did not observe a correlation between cell survival fractions and RSG or ATP levels (**Fig. 3e, f**). This lack of correlation may stem from the multifaceted nature of bacterial survival strategies, which should involve various cellular processes beyond metabolic activity.

### Whole genome sequencing revealed diverse genetic attributes

High antibiotic tolerance and resistance should be attributed to the accumulation of mutations; therefore, we conducted comprehensive whole-genome sequencing for each sample (see Materials and Methods). The sequencing data revealed three distinct types of mutations: insertions and deletions (INDELs), structural variations (SVs), and single nucleotide polymorphisms (SNPs) (Supplementary **Table S2**). Although mutations in the same genes were occasionally observed across the samples, there was no evidence that the evolution of resistant mutants in these samples converged on a common pathway or mechanism (**Fig. 4**). The Venn diagram illustrates the distribution of mutations within the samples, highlighting that only a select few genes, such as *cyoE* (present in S1 and S5), *rhsC* (found in S2 and S6), and *adk* (shared by S5, S6, and S7), exhibited these common mutations within their genomes (**Fig. 4**). However, we noted SNP mutations in one of the metabolic genes, *icd*, which were present in most samples except S4 and S6 (Supplementary **Table S2**), aligning with existing literature [21].

**Fig. 4.**
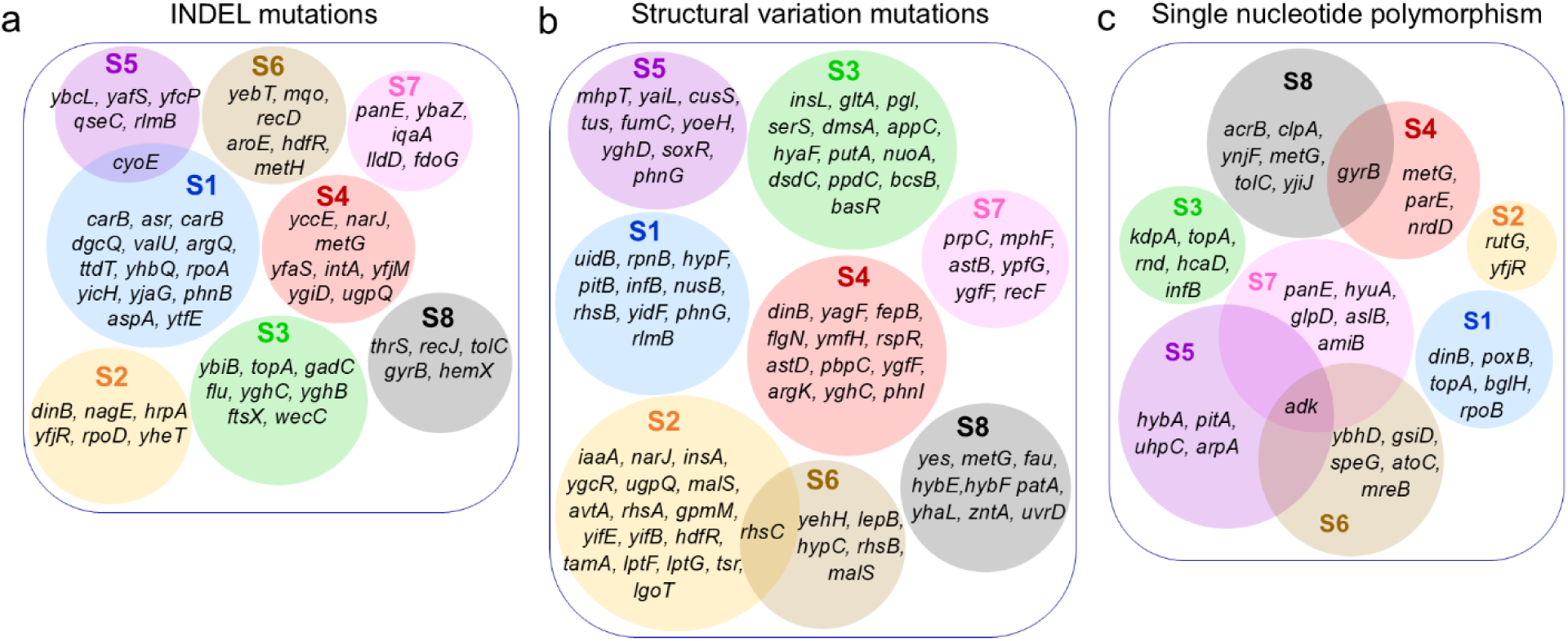
Whole genome sequencing reveals that various types of mutations have occurred within the evolved strains. A Venn diagram illustrates the mutations identified in the mutants as a result of the adaptive laboratory evolution experiment, including **(a)** insertions and deletions (INDELs), **(b)** structural variations (SVs), and **(c)** single nucleotide polymorphisms (SNPs).

To gain more insights into how the mutations in the identified genes impact cellular antibiotic resistance, tolerance, and fitness parameters, we individually deleted 50 genes (including those exhibiting the highest frequency of mutations among the evolved samples) in the *E. coli* MG1655 strain. Given that each sample contains numerous mutations, including SNPs, INDELs, and SVs across a wide range of genes, the combinatorial effects of these mutations make it nearly impossible to gain mechanistic insights for each strain. However, creating single-gene deletions for some of the observed mutations and analyzing their survival fractions, MIC levels, and fitness parameters could help us understand the fluctuations in tolerance, resistance, and fitness traits observed in the evolved samples. Therefore, we performed the aforementioned assays, mirroring those conducted on the evolved mutants, for the knockout strains we generated. Since our background *E. coli* MG1655 strain, used for gene deletions, did not have the mCherry expression cassette, we could not determine competition or non-growing cell scores. However, we were still able to determine the critical fitness parameters such as doubling times, lag scores, and other metabolic properties (Supplementary **Fig. S4** and **Fig. S5)**. In the case of survival fraction, approximately 25-26 single knockout strains (some of them are Δ*icd*, Δ*cyoE*, Δ*lgoT*, Δ*yghC*, Δ*tolC*, Δ*rnd*, Δ*dld*, Δ*acrB*, Δ*ybcL*, Δ*uidB*, Δ*dsdC*, Δ*narJ*, Δ*yoeH*, Δ*recF*, Δ*rhsC*, Δ*dinB*, Δ*flgN*, Δ*hybE*, Δ*yjeP*, and Δ*yghD*) displayed enhanced ofloxacin tolerance compared to the wild-type control strain (**Fig. 5a**). A smaller set of strains (Δ*ygfF*, Δ*adk*, Δ*gpmM*, Δ*gltA*, Δ*fau*, Δ*nuoA*, and Δ*uvrD*) exhibited notably reduced ofloxacin tolerance in contrast to the wild-type control (**Fig. 5a**). Several mutants, including Δ*tolC*, Δ*acrB*, and Δ*uvrD*, exhibited significantly lower MIC levels, whereas Δ*nuoA* exhibited slightly higher MIC levels compared to the wild type (**Fig. 5b**). However, no correlation was observed between the MIC levels and the survival fractions of these mutant strains (**Fig. 6a**). Simultaneous mutations that may increase or decrease tolerance or resistance could explain the observed fluctuations in survival rates and MICs of the mutant strains.

**Fig. 5.**
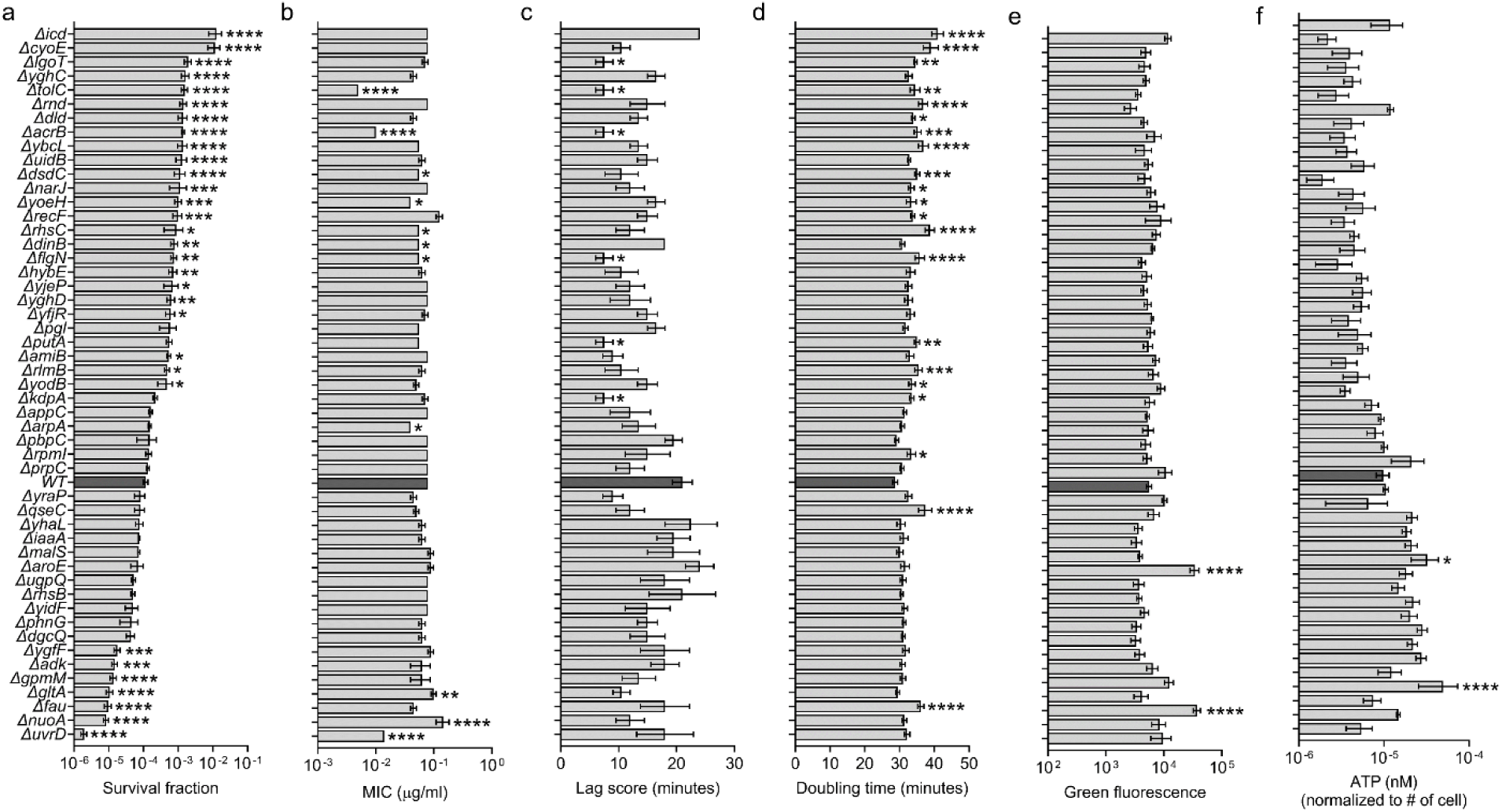
Survival fraction, MIC, fitness factors, and metabolic activities of the single mutants revealed new genetic determinants for tolerance. **(a)** Stationary phase cells of the designated individual single mutants were exposed to ofloxacin (5 µg/ml) after diluting 100-fold in LB media for 7 h. Treated cells were collected, washed, and plated on LB agar plates to enumerate the CFU levels. n=4. **(b)** Indicated single mutants were diluted to have ∼10^8^-10^9^ cells/ml and spread on a circular LB agar plate. Agar plates were dried for 20 minutes next to the flame. Ofloxacin ETEST strips were placed on the dried agar plates and incubated for 16 h to determine the minimum inhibitory concentrations. n=4. **(c)** Lag scores of the indicated individual mutant strains were calculated using the growth curves (Supplementary **Fig. S4**). n=4. **(d)** Doubling times of the indicated individual strains were calculated using the exponential phase of the growth curves (Supplementary **Fig. S4**). n=4. **(e)** Stationary phase cells of individual single mutants were stained with RSG dye and analyzed with a flow cytometer to determine the metabolic state of the cells (Supplementary **Fig. S5**). n=4. **(f)** Stationary phase cells of the indicated strains were collected to measure the intracellular ATP concentrations (see Materials and Methods). A flow cytometer was used to count the cell number for normalization purpose. n=4. Statistical analysis was performed between the WT and single mutants using one-way ANOVA with Dunnett’s post-test. *P<0.05, **P < 0.01, ***P < 0.001, and ****P < 0.0001. Data corresponding to each time point represents mean value ± standard deviation.

**Fig. 6.**
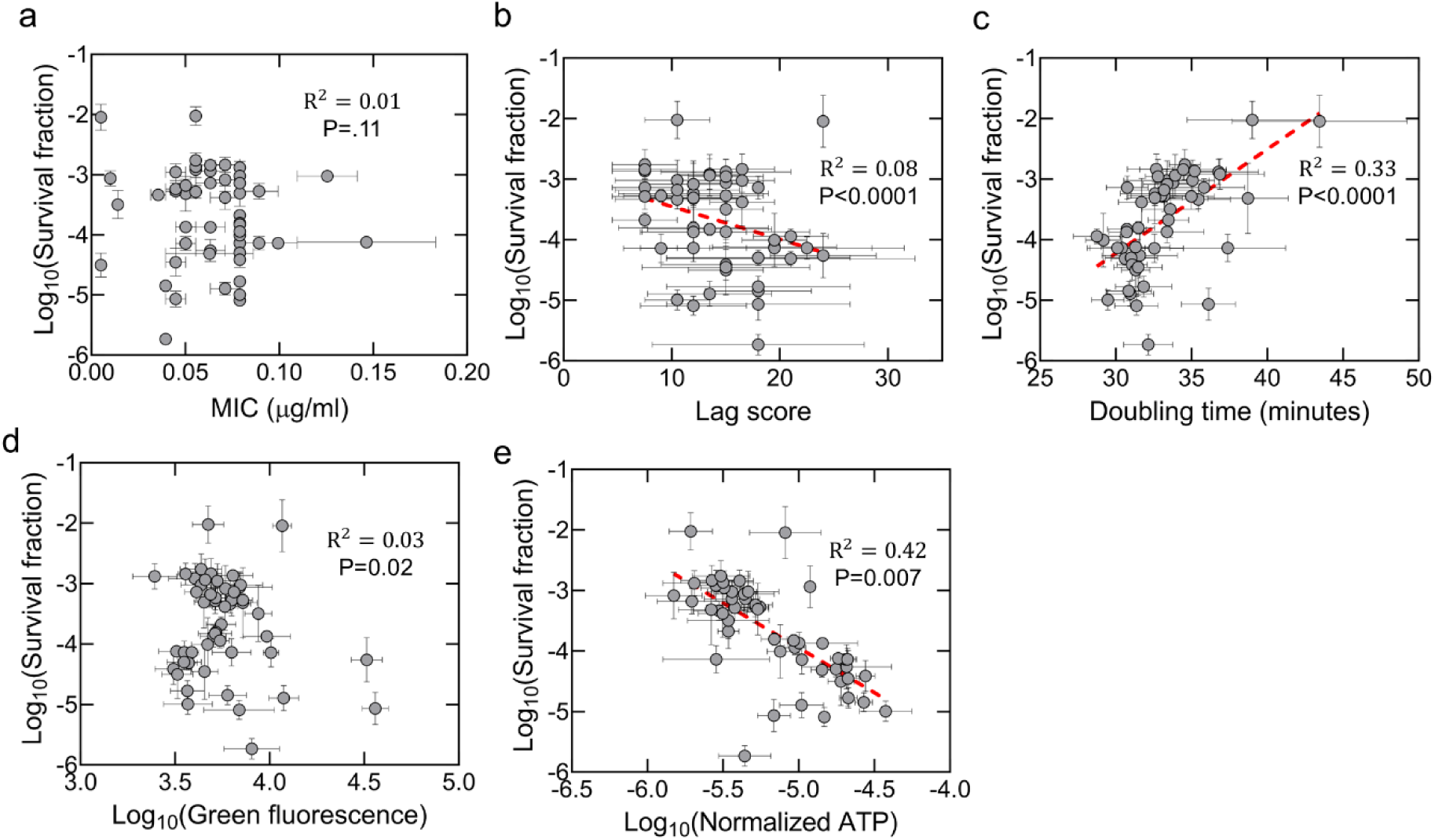
Correlation between survival fractions, fitness factors, and metabolic parameters of knockout strains. **(a)** The survival fraction and MIC of the single knockout strains were plotted, and a linear regression analysis was performed for the correlation analysis (data from **Fig. 5b**). **(b)** Correlation between survival fractions and lag scores. A linear regression analysis was performed between survival fractions and lag scores of the single knockout strains (data from **Fig. 5c**). **(c)** Correlation between survival fractions and doubling times. A linear regression analysis was performed between survival fractions and doubling times of the single knockout strains (data from **Fig. 5d**). **(d)** Correlation between survival fractions and cellular redox states. A linear regression analysis was performed between survival fractions and metabolic states of the single knockout strains (data from **Fig. 5e)**. **(e)** Correlation between survival fractions and ATP levels. A linear regression analysis was performed between survival fractions and ATP levels of the single knockout strains (data from **Fig. 5f**). Statistical significance was determined using F-statistics. Data corresponding to each time point represents mean value ± standard error.

When we analyzed the fitness factors of the generated knockout strains, we found that strains with higher survival fractions tended to have slightly higher doubling time and lower ATP values (**Fig. 5d, f**), and observed a moderate correlation between survival fractions and these two parameters (**Fig. 6c, e**). While we did not detect any clear trend or correlation between survival fractions and the redox levels of the strains (**Fig. 5e** and **Fig. 6d**), interestingly the strains with higher tolerance exhibited slightly lower lag scores (**Fig. 5c** and **Fig. 6b**). Nevertheless, our data clearly show that knockout strains exhibit diverse fitness characteristics irrespective of their tolerance or resistance levels (**Fig. 5a-f**), and correlations between fitness and survival are not always strong or consistent (**Fig. 3** and **Fig. 6**).

## Discussion

ALE experiments provide us valuable insights into how bacteria develop resistance, especially under the rapid evolutionary pressures of antibiotic treatment [16,17,19,21,46]. In this study, out of the eight *E. coli* samples from the ALE experiments, seven samples, S1 and S3-S8, showed better survival characteristics during ofloxacin treatments compared to the control group. The fact that S2 did not show improved survival over 22 days reveals the unpredictability of evolutionary processes and how results can vary even in well-controlled experiments. This variation might be due to factors like the type and frequency of mutations and microscopic-scale environmental changes [60,61]. Our findings also revealed that there is no correlation between survival rates and MIC levels, suggesting that tolerance and resistance to ofloxacin might evolve independently in cyclic ALE experiments. When faced with selective pressure, bacteria can adopt various survival strategies, leading to a wide range of phenotypic variants, with some managing to survive high doses of antibiotics without necessarily becoming more resistant.

We demonstrated that the ten randomly chosen colonies from each sample consistently had similar survival rates and MIC levels. This suggests that exposing the cells to antibiotics repeatedly enriched mutant strains with consistent traits within each sample. This uniformity might come from a few mutant lineages exhibiting greater fitness and survival characteristics; the cyclic pressure from ofloxacin during the ALE process may favor these lineages, eventually leading to the dominance of these specific traits within each sample. However, despite this consistency within individual sample, there was significant diversity across the eight samples. It is obvious that different mutant lineages within each sample followed unique evolutionary paths. Even though all samples started from the same strain, random mutations during the ALE process could have created genetic diversity among the samples. These genetic differences should explain the varying responses to ofloxacin, resulting in different survival rates and MIC levels across samples.

We explored fitness traits of the samples (S1-S8) to establish if there is a trade-off between antibiotic tolerance/resistance and these traits. Except for S2, all samples had slightly longer doubling times than the control. In co-culture environments, samples S4 and S8 showed notable growth suppression, indicating a decreased capacity to compete with parental cells. There seems to be a connection between survival and competitive advantage, as evidenced by the moderate correlation found between survival fractions and doubling times or competition scores of the samples. However, we found no evidence of a correlation between bacterial survival and other fitness parameters linked to non-growing cell counts, lag scores, and metabolic processes. Previous studies have identified the lag phase as a critical factor in the evolution of antibiotic-resistant strains [16,48], and traditional models suggest that resistance mutations often impose fitness costs [49,50,62]. While we fully agree with these perspectives, it is important to emphasize that these traits are contingent on the environmental conditions that organisms encounter. For instance, cyclic ALE experiments might select for mutations that confer resistance with minimal fitness trade-offs, as the recovery phase allows resistant mutants to compete and proliferate. This process can lead to the selection of optimized resistance mechanisms, favoring mutations that do not drastically impact metabolic efficiency or growth capabilities. The alternating selective pressures of cyclic ALE mimic real-life fluctuating environments, and bacteria evolving under these conditions may develop more robust and adaptable resistance mechanisms that perform well under both stress and non-stress conditions.

We carried out whole genome sequencing to get further insight into genetic backgrounds of the samples that have diverse fitness traits and antibiotic resistance and tolerance properties. We don’t think that the evolution of resistance appears to follow a consistent pathway or mechanism, even though a few mutations in the same genes occur in different samples. Common mutations found in samples for particular genes, such *icd, cyoE*, *rhsC*, and *adk*, representing potential targets for further investigation. We are aware that each sample has diverse mutations in numerous genes, and these mutations may eliminate, enhance, or completely change the functions of these genes. Given the combinatorial impact of all these mutations on antibiotic tolerance and resistance, it is very challenging to gain a mechanistic understanding from these diverse mutational landscapes. Nevertheless, we deleted individual genes, including those exhibiting the highest frequency of mutations, to assess their effects on cellular antibiotic resistance, tolerance, and fitness characteristics. When compared to wild-type strains, a significant number of knockout strains exhibit increased ofloxacin tolerance, as expected. While our research supports the prior findings that several of these knockout strains, including metabolic mutations, enhance bacterial survival [21], we also found new genes potentially associated with antibiotic tolerance and/or resistance, including Δ*cyoE,* Δ*lgoT,* Δ*yghC,* Δ*rnd,* Δ*udiB,* Δ*dsdC,* Δ*narJ,* Δ*yoeH,* Δ*flgN,* Δ*hybE,* Δ*yjeP,* Δ*yghD,* Δ*yfjR,* Δ*amiB,* Δ*adk,* and Δ*ygfF*. The roles of these genes remain unclear, offering promising directions for future research into their contributions to antibiotic resistance and tolerance. Additionally, we observed few knockout strains Δ*uvrD,* Δ*nuoA,* and Δ*gltA,* with decreased tolerance, which is consistent with previous studies [63,64]. The concurrent emergence of mutations that may either enhance or reduce tolerance could account for the observed fluctuations in the survival fractions of the mutant strains.

Many fitness parameters of the knockout strains do not show a clear trend or correlation with survival fractions although some knockout strains with greater survival fractions tend to have slightly higher doubling scores and lower ATP values. The observed variations in metabolic activities among the knockout strains may indicate that bacterial cells may use diverse metabolic strategies to deal with antibiotic-induced stress. In fact, our whole genome sequencing revealed the enrichment of mutations in several metabolic genes, such as *icd*, *nuo,* and *gltA*. This is in line with recent research that highlights the possible contribution of metabolic adaptations to antibiotic tolerance. Henimann’s and Michiels’ groups investigated genomic alterations in *E. coli* strains— including uropathogenic UTI89—that were exposed to antibiotics periodically and observed mutations in the genes of the *nuo* operon, a critical element of *E. coli* energy metabolism [65]. Additionally, the work conducted by Collin’s group identified hitherto unidentified metabolic genes (*icd, gltD,*and *sucA*) that are implicated in the development of antibiotic resistance in *E. coli* cells exposed to a variety of antibiotics [21]. They further supported these results by analyzing a huge library of 7243 *E. coli* genomes from NCBI Pathogen Detection, showing that comparable mutations are present in clinical *E. coli* infections [21].

The deletion of the *icd* gene has been repeatedly shown to increase the antibiotic tolerance of bacterial cells by various studies [21,66,67]. Although the identification of mutations in the *icd* gene in most samples, including S2, in our study might imply a conserved evolutionary mechanism, the mutations in this gene alone do not fully explain the observed tolerance, as S2 does not exhibit increased tolerance or resistance. This may imply the collective effects of the multiple mutations in shaping the evolutionary pathways of resistant or tolerant strains. Additionally, our data show that not all metabolic gene knockout strains carry significant fitness costs, while the *icd* knockout exhibit such costs. This is also supported by several recent studies [66,67]. This phenomenon could give a survival advantage to bacterial cells, as they may acquire metabolic mutations to increase their tolerance without experiencing significant fitness costs.

In this study, we used ofloxacin as the antibiotic for the ALE experiment, which typically induces mutations in target genes like *gyrA, gyrB, parC,* and *parE*. Previous studies have observed such mutations when antibiotics were continuously present in cultures, with concentrations gradually increased to drive resistance evolution [68–70]. While we observed few mutations in the drug targets (except *gyrB*), this could be due to our treatment approach, where cells were exposed to high concentrations of ofloxacin and then transferred to fresh medium without antibiotics, allowing the surviving cells to compete. Since most antibiotic target proteins are essential and involved in cell growth, mutations in these proteins may significantly reduce cell fitness [71,72], making them less competitive during recovery.

In conclusion, in this study, we demonstrated how bacterial cells acquire complex resistance or tolerance mechanisms to survive unfavorable conditions. Although we observed a few common mutations across different evolved samples, we did not find a clear pattern of evolution leading to a conserved pathway or mechanism. Our study, which uncovers the complex relationship between antibiotic tolerance, genetic mutations, and bacterial fitness, adds to our understanding of bacterial adaptation mechanisms, which may help guide the development of therapeutic interventions to address the heterogeneous nature of antibiotic resistance.

## Materials and Methods

### Bacterial strains and plasmids

In this study, we used *Escherichia coli* K-12 MG1655 and MO strain. Unless noted otherwise, all the knockout strains in *E. coli* MG1655 were generated during the study. *E. coli* MO strain harbors red fluorescence (*mCherry* gene) expression system in the genome [55,58]. pUA66-*gfp* expression system was used with kanamycin-resistant gene as selection marker. Both *mCherry* and *gfp* are isopropyl β-d-1-thiogalactopyranoside (IPTG) inducible and tightly regulated by *T5* and *LacI^q^* as promoter and repressor, respectively. A complete description of bacterial strains and plasmids used in this study is listed in Supplementary **Table S1**. Primers used to generate knockout strains are tabulated in Supplementary **Table S3** and **Table S4**. *mCherry* and *gfp* expressions were used to monitor the cell proliferation at the single-cell level with flow-cytometry. Fluorescent protein overexpression through chromosomally integrated *mCherry* and plasmid pUA66-*gfp* did not alter the antibiotic sensitivity of *E. coli* cells. An empty vector of pUA66 plasmid (without *gfp* gene) was used as a control throughout the study.

### Media, chemicals, and culture conditions

Unless otherwise noted, chemicals utilized in this study were purchased from Fisher Scientific (Atlanta, GA), VWR International (Radnor, PA), or Sigma-Aldrich (St. Louis, MO). To grow the bacteria in liquid media, Lysogeny Broth (LB) was prepared from its components which include yeast extracts (5 g per 1 L DI water), tryptone (10 g per 1 L DI water) and sodium chloride (10 g per 1 L DI water). LB agar (40 g premixed per 1 L DI water) was used as solid media to grow the bacteria whenever required. IPTG (1 mM as final concentration) was used to overexpress fluorescent proteins (mCherry or GFP) in *E. coli* [55,73]. For plasmid retention and antibiotic selection marker, 50 µg/ml kanamycin was used. To perform *in vitro* clonogenic survival assay on *E. coli* cells, 5 µg/ml ofloxacin was used. Phosphate Buffered Saline solution was used to remove chemicals and antibiotics from the liquid cell cultures before spotting them on the agar plate to enumerate the surviving cells. Ultra-pure distilled water was used to prepare chemical and antibiotic solutions as well as growth media. NaOH solution was used to dissolve ofloxacin in water, where the final concentration of NaOH became 0.06 mM. Bacterial growth media was sterilized by autoclaving at high temperature (121°C) and pressure (15 psi). Chemical and antibiotic solutions dissolved in water were sterilized using a 0.2 µm polyether sulfone membrane filter. FlowJo Software (Tree Star Inc., Ashland, OR) was used to analyze all the flow cytometry samples in this study. Overnight pre-cultures were prepared by inoculating 2 ml LB media in 14-ml falcon tubes from a frozen (-80 ⁰C) cells stock and cultured at 37 ⁰C with shaking (250 rpm) for 16 h.

### Adaptive laboratory evolution technique

To enrich the *E. coli* cultures with tolerant or resistant cells, the adaptive laboratory evolution (ALE) technique was adopted. Overnight pre-cultures of *E. coli* MO strain harboring pUA66-empty vector (EV) were diluted 100-fold (∼3-5×10^7^ cells/ml) in 2 ml LB media in a 14-ml falcon tube followed by the treatment with 5 µg/ml of ofloxacin (OFX) and cultured at 37 ⁰C with shaking (250 rpm). For plasmid maintenance, kanamycin (50 µg/ml) was added to the LB media [73]. At the end of the 7-h treatment, 1 ml treated cells were washed three times with PBS (1X) by centrifugation (13,300 rpm or 17,000 xg) to remove antibiotics. After the final washing step, pelleted cells were inoculated in 2 ml LB media and grown in the shaker as an overnight pre-culture for the subsequent day. To determine the number of survived cells after antibiotic treatment, the remaining 1 ml of treated cells were washed as described above. At the final washing step, 900 µl supernatant was removed, and pelleted cells were resuspended in the remaining 100 µl PBS. Resuspended cells were serially diluted in 90 µl PBS. Ten microliter cell suspension was spotted on the agar plate to enumerate the survived cells. Before ofloxacin treatment, 10 µl cell suspensions were serially diluted in 90 µl PBS and spotted on the agar plate to count the initial number of cells. Agar plates with cells were incubated at 37 ⁰C for 16 h to allow *E. coli* colony formation. This cyclic intermittent antibiotic treatment was carried out for 22 days without interruption. After 22 days, 25% glycerol cell stocks were prepared for the evolved cell samples and stored at -80 ⁰C for future assays.

To confirm the homogeneity of ofloxacin resistance/tolerance among the individual evolved cell samples, frozen cell samples were streaked on LB agar plates containing 50 µg/ml kanamycin (to avoid potential contamination). Agar plates were incubated at 37 ⁰C for 16 h. Ten random colonies from each evolved cell sample were picked and cultured in LB media in a 96-well plate format at 37 ⁰C with shaking. Breath-easy Membrane was used to cover the 96-well plate. Once the cultures reached the stationary phase (t=16 h), 25% glycerol cell stock of each culture was prepared and stored at -80 ⁰C for subsequent assays.

### Cell growth assay and calculations of lag and doubling time scores

Compared to the parental strain, cell growth was measured in optical density to assess the growth trend of the eight evolved samples. The growing cultures’ optical density (OD_600_) was monitored at 600 nm wavelength using a plate reader (Varioskan LUX Multimode Microplate Reader, Thermo Fisher, Waltham, MA, United States) for 24 h. Cell growth assay cultures were prepared by diluting overnight pre-cultures 100-fold (∼3-5×10^7^ cells/ml) in 2 ml LB media in 14-ml falcon tubes and grown at 37 ⁰C with shaking (250 rpm). At designated time points 300 µl cell suspension was placed in the 96-well flat bottom plates and measured the OD_600_ with the plate reader. The lag score, representing the time bacteria take to initiate growth in a specific medium, was calculated using time-dependent growth data and determined based on the methodology outlined in previous studies [56,57]. Doubling time is the duration it takes for bacteria to double their population size through binary fission. We calculated the doubling time using data from the exponential growth phase with the following formula:

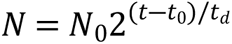

where, N_0_ is the initial number of cells when the exponential phase starts at t_0_, N is the number of cells at time t, and t_d_ is the doubling time.

### Antibiotic tolerance assay

Overnight pre-cultures were diluted 100-fold (∼3-5×10^7^ cells/ml) in 2 ml LB media in a 14-ml falcon tube and immediately treated with antibiotics (ampicillin, gentamycin, and ofloxacin) and cultured at 37 ⁰C with shaking (250 rpm). Kanamycin (50 µg/ml) was added to the culture to retain the plasmid. To determine the initial number of cells before antibiotic treatment, 10 µl cell suspension was serially diluted in 90 µl PBS and spotted on the agar plate. At the designated time point after treatment (i.e., 7-h treatment), 1 ml of treated cells were washed twice with PBS (1X) to reduce the antibiotic concentration at the sub-MIC level by centrifugation (13,300 rpm or 17,000 xg). After the final centrifugation, 900 µl supernatant was removed, and pelleted cells were resuspended in the remaining 100 µl PBS. Resuspended cells were serially diluted in 90 µl PBS to be able to count once plated in an agar plate. Then, serially diluted cell suspensions were spotted on the agar plates to enumerate the colony formation units (CFU), as described above. To grow the *E. coli* colonies, agar plates were incubated at 37 ⁰C for 16 h.

### Non-growing cells detection assay

To determine the level of non-growing cells in the evolved samples and the parental strain (*E. coli* MG1655 MO strain harboring pUA66-empty vector), the IPTG-inducible red fluorescent protein (mCherry) expression system was used. To obtain the red fluorescence protein (mCherry) positive cells, 1 mM IPTG was added in the overnight pre-cultures. mCherry-positive cells (1 ml) from the overnight pre-cultures were washed with PBS three times to remove IPTG and resuspended in 1 ml sterile LB media. Washed cells were diluted 100-fold in 2 ml LB broth without the inducer (IPTG) in 14-ml falcon tubes and cultured at 37 ⁰C with shaking (250 rpm). At designated time points (t=0, 1, 2, and 3 h), cells were diluted in PBS and their red fluorescent protein (mCherry) levels were measured through a flow cytometer (NovoCyte Flow Cytometer, NovoCyte 3000RYB, ACEA Biosciences Inc., San Diego, CA, United States). We note that cells were always diluted in PBS to achieve a cell density (∼10^6^-10^7^ cells/ml) for flow analysis. At the beginning of the cell culturing without inducer (t=0), cells expressed high levels of red fluorescence. Later, once cells started to proliferate, due to the protein dilution, cells gradually lost their red fluorescence protein levels. Finally, at t=3 h, a small subpopulation of the culture, consisting of non-growing cells, still exhibited high levels of red fluorescent protein due to a lack of cell division. The fractions of these non-growing cells were determined by flow cytometry. The excitation wavelength for red fluorescence was 561 nm, and a 615/20 bandpass filter received the fluorescence signals.

### Competition assay

For the competition assay, equal numbers of cells from both the parental strain (*E. coli* MG1655 MO strain harboring pUA66-*gfp*) and an evolved strain (S1, S2, S3, S4, S5, S6, S7, and S8) were diluted in 2 mL of LB media from overnight pre-cultures and grown at 37°C with shaking (250 rpm) for 24 h. To induce red (mCherry) and green (GFP) fluorescent protein, 1 mM IPTG was added in the pre-cultures and competition cultures. We note that parental strain harboring plasmid expressing GFP was used to be able to monitor their growth through a flow cytometer. During the co-culturing, at designated time points (t= 0, 4, 8, and 24 h), cells were collected and diluted in PBS to analyze through a flow cytometer. Cells were diluted in PBS to reach a cell density of around 10^6^-10^7^ cells/ml for the flow cytometry analysis. The excitation wavelength for green fluorescence was 488 nm and a 530/30 bandpass filter collected signals. The excitation wavelength for red fluorescence was 561 nm, and a 615/20 bandpass filter received the fluorescence signals. To maintain the plasmid, kanamycin (50 µg/ml) was added to the LB medium.

### Minimum inhibitory concentrations (MICs) determination

Overnight pre-cultures of parental strain and evolved cells were prepared as described above. When necessary, kanamycin (50 µg/ml) was added to the culture to retain the plasmid. Agar plates were prepared using circular Petri dishes (100×15 mm, Fisher Scientific, catalog# FB0875712) into which approximately 15–20 ml of melted LB agar was poured. Overnight pre-cultures of each sample were diluted (∼5×10^8^ cells/ml) in 1 ml PBS and spread with an L-shape sterile spreader on the agar plate. Inoculated plates were dried at room temperature (∼25 ⁰C) next to a flame for 15 minutes. Ofloxacin (OFX) MIC Test Strip (Fisher Scientific, catalog# 22-777-876) was placed in the dried agar plate and incubated at 37 ⁰C for 20 h. We note that the MIC Test Strip contained known concentration gradients of ofloxacin ranging from 0.002 to 32 µg/ml.

### Next generation sequencing

To identify the genetic perturbations on the genome of the evolved cells, short-read non-human whole genome sequencing (next-generation sequencing) was performed in GeneWiz (South Plainfield, NJ, USA). Overnight pre-cultures of the parental and evolved strains were diluted 100-fold (∼3-5×10^7^ cells/ml) in 2 ml LB media with 50 µg/ml kanamycin in 14-ml falcon tubes and grown at 37 ⁰C with shaking (250 rpm). Late-exponential phase (OD_600_∼1.0) cells were collected and washed twice with PBS (1X) by centrifugation (13,300 rpm or 17,000 xg). At least 100 µl cell pellet was collected in Eppendorf tubes after the final washing step. Eppendorf tubes containing pelleted cells of the individual sample were immediately immersed in a dry ice-ethanol bath (∼-80 ⁰C) for 10 minutes. Frozen cell pellets were stored at -80°C before being shipped to GenWiz. Notably, the wild-type parental strain was used as the background genome sequence control. The sequencing details from GeneWiz can be found elsewhere [74].

### Statistical analysis

At least four biological replicates were performed for each condition except the generation of evolved cells. To assess the statistical significance between control and treatment groups, a one-way ANOVA test with Dunnett’s posttest was performed where the threshold value of P was chosen as * P<0.05, ** P<0.001, *** P<0.0001. Average ± Standard Deviation was used to present all the data points in the linear graphs. In the case of flow diagrams, a representative figure from among all the replicates is presented in this article, while other replicates showed a similar trend.

## Acknowledgments

The authors would like to thank the members of Orman Lab for their help.

## Author contributions

S.G.M. and M.A.O. conceived and designed the study. S.G.M., P.K., D.F., and S.G. performed the experiments. S.G.M. and M.A.O. analyzed the data and wrote the paper. All authors have read and approved the manuscript.

## Data and materials availability

All relevant data are provided in the main figures or supplementary files. The raw data have been deposited in FigShare https://doi.org/10.6084/m9.figshare.27935040.v1 [75], and the genomic sequencing data have been submitted to ArrayExpress and will be made available during the review process upon receipt of the accession number.

